# Atomic structure of human sapovirus capsid by single particle cryo-electron microscopy

**DOI:** 10.1101/2021.02.09.429995

**Authors:** Naoyuki Miyazaki, Kosuke Murakami, Tomoichiro Oka, Motohiro Miki, Kenji Iwasaki, Kazuhiko Katayama, Kazuyoshi Murata

## Abstract

Sapovirus is a cause of acute gastroenteritis in humans and animals. Infants and younger children have the greatest disease burden. Although it shares many similarities with norovirus, the lack of detailed structural information has hampered the development of vaccines and therapeutics. Here, we investigated the human sapovirus VLP by single particle cryo-electron microscopy and are the first to report the atomic structure of the capsid at 2.9 Å resolution. The atomic model revealed the domain interactions of the capsid protein and functionally important amino acid residues. The extended loop from the P1 subdomain was involved in interactions in the P2 domain, forming unique arch-like dimeric protrusions of capsid proteins. All hypervariable regions that are important candidates for immune response or receptor binding, formed a large cluster at the top of the P domain. These results pave the way for developing vaccines, antiviral drugs, and diagnostic systems for this infectious disease.

## Introduction

Sapovirus (SaV) belongs to the *Caliciviridae* family and is well known to cause acute gastroenteritis in humans as well as animals. Human SaV (HuSaV) contains a positive-sense single-stranded RNA genome of approximately 7.1 to 7.5 kb in length (Green et al., 2001) and is divided into four genogroups (GI, GII, GVI, and GV), although animal SaVs are further diverged and divided into 19 genogroups (Farkas et al., 2004; Yinda et al. 2017; Li et al., 2018). The viral genome consists of two or three open reading frames (ORFs). ORF1 encodes a polyprotein that undergoes proteolytic cleavage to form non-structural proteins and a major capsid protein VP1 (viral protein 1). The major capsid protein VP1 is solely responsible for most capsid-related functions, such as assembly, host interactions, and immunogenicity. ORF2 encodes a minor structural protein, VP2. In the case of a feline calicivirus (FCV), a member of genus *Vesivirus*, the minor structural protein VP2 forms a large dodecameric portal-like assembly at a unique three-fold axis of icosahedral symmetry after receptor engagement (Conley et al., 2019), which likely functions as a channel for genome release from the capsid. ORF3 encodes a small basic protein of unknown function (Clarke and Lambden, 2000; Atmar and Estes, 2001). Compared with well-characterized norovirus and vesivirus, there are limited studies on SaV. In fact, the SaV structure has been reported only at low and intermediate resolutions (Chen et al., 2004; Miyazaki et al., 2016), and the atomic structure of the SaV capsid has not yet been elucidated. The SaV belongs to a different genus from the well-characterized caliciviruses and has different host specificity and immunogenicity. In addition, an understanding of replication strategies, pathogenesis, and immunogenicity of SaV have also been hampered due to the lack of a sufficient viral replication system, such as the actual target cells in the host, until the recent establishment of the SaV cultivation system (Takagi et al., 2020).

The *Caliciviridae* family is currently classified into eleven established genera: *Bavovirus, Lagovirus, Minovirus, Nacovirus, Nebovirus, Norovirus, Recovirus, Salovirus, Sapovirus, Valovirus*, and *Vesivirus* (Vinjé et al., 2019). The atomic structures of calicivirus VLPs or virions in three established genera have been determined for Norwalk virus (NV; in genus *Norovirus*), rabbit hemorrhagic disease virus (RHDV; in genus *Lagovirus*), San Miguel sea lion virus (SMSV; in genus *Vesivirus*), and FCV (in genus *Vesivirus*), while atomic structures in other genera, including *Sapovirus*, remain unknown (Prasad et al., 1999; Chen et al., 2006; Ossiboff et al., 2010; Wang et al., 2013; Song et al., 2020). The calicivirus virions have a mostly conserved capsid shell, which is composed of 180 copies of VP1 arranged in a T=3 icosahedral symmetry. The VP1 proteins are designated A, B, and C in the icosahedral asymmetric unit according to their positions, which form quasi-equivalent A/B and C/C dimers (Prasad et al., 1999; Chen et al., 2006; Ossiboff et al., 2010). Each VP1 capsid monomer contains two principal domains, shell (S) and protrusion (P) domains, with an N-terminal arm located inside the capsid shell. The S domains represent the most conserved region of the amino acid sequence among caliciviruses, and have a classical eight-stranded β sandwich, which has been commonly found in T=3 icosahedral viruses (Rossmann and Johnson, 1989). The fundamental function of the S domains is to form a contiguous icosahedral capsid shell responsible for the protection of their viral genome from the outer environment. In contrast, structures and amino acid sequences of the P domain are rather variable among caliciviruses because this domain is involved in virus-host interactions and immunogenicity. Indeed, the sizes of the P2 domains are considerably different between *Norovirus* and *Vesivirus*, which are 127 and 176 amino acid residues, respectively (Prasad et al., 1999; Chen et al., 2006). The P domains form protrusions on the capsid shell composed of the S domains and each P domain can be further divided into two subdomains called P1 and P2. In spite of the little sequence conservation, protein folds of the P1 and P2 subdomains are conserved among caliciviruses, but unique to other viruses except for hepatitis E virus (HEV: Yamashita et al., 2009; Guu et al., 2009; Xing et al., 2010). The relative orientation between S-P1-P2 domains shows inter-genus variations (rarely including intra-genus variations). For example, only the P2 domain is involved in the dimeric interactions in *Vesivirus*, while both the P1 and P2 domains participate in the dimeric interactions in *Norovirus*. Because of the inter-genus diversities, we are eager to elucidate the atomic structures of other genera, including *Sapovirus*, in order to understand their immunogenicity and virus-host interactions in more detail.

Here, we determined the capsid structure of a HuSaV virus-like particle (HuSaV-VLP) at 2.9 Å resolution by single particle cryo-electron microscopy (cryo-EM), and successfully built an atomic model of the capsid. The atomic model revealed the domain boundary and the functionally important amino acid residues in the capsid protein, 1) the unique arch-like dimeric protrusion on the capsid surface provides hints for a stable construct design for vaccine development, and 2) the detailed structure of the large hypervariable region cluster at the top of the P domain accelerates the development of vaccines and antivirals.

## Results and Discussion

### Structure determination of HuSaV VLP

The capsid protein VP1 of HuSaV from the Nichinan strain in genogroup I (GI. Nichinan; Iwakiri et al., 2009) was expressed in a baculovirus expression system, and the self-assembled and secreted HuSaV-VLPs from the cells were purified from the culture medium. The three-dimensional (3D) structure was determined at 2.9 Å resolution by cryo-EM single-particle analysis (Figures 1, S1, and S2). The cryo-EM map clearly shows a T=3 icosahedral symmetry with 90 protrusions, composed of 180 copies of the VP1 protein in total, distributed along the icosahedral 2-fold axes (C/C dimers) and quasi 2-fold axes (A/B dimers) on the surface (Figures 1A and S2A). The bulky side chains are clearly resolved in the cryo-EM, and the atomic models are unambiguously built for the VP1 proteins (Figures 1B and S1E). The cryo-EM map allowed atomic modeling of residues 38–554 for subunit A, residues 21–554 for subunit B, and residues 21–554 for subunit C, except for one disordered loop (residues 380–384) for all subunits (Figure 2). The icosahedrally independent A/B and C/C dimers show slightly different conformations, mainly between S-domains in each dimer, which are the bent and flat conformations, respectively, as in other T=3 viruses (Figure S3). The βI-βA’ loop between the S-domain and P-domain, containing a β-turn (residues 231–234), undergoes a hinge-like motion to adapt the two conformers (red asterisks in Figure S3). In addition, N-terminal regions of residues 21–37 in the B- and C-subunits extend underneath the capsid shell and interact with the neighboring subunits around icosahedral 3-fold axes (Figure 3A), although the N-terminal region is disordered in the A-subunit, and no N-terminal network is observed around icosahedral 5-fold axes. In particular, residues A25-T26 form a short inter-subunit β-sheet with F154-V155 in the adjacent subunit (Figure 3B). Therefore, the N-terminal network around the icosahedral 3-fold axes likely stabilized the hexameric units in the capsid.

**Figure 1.**
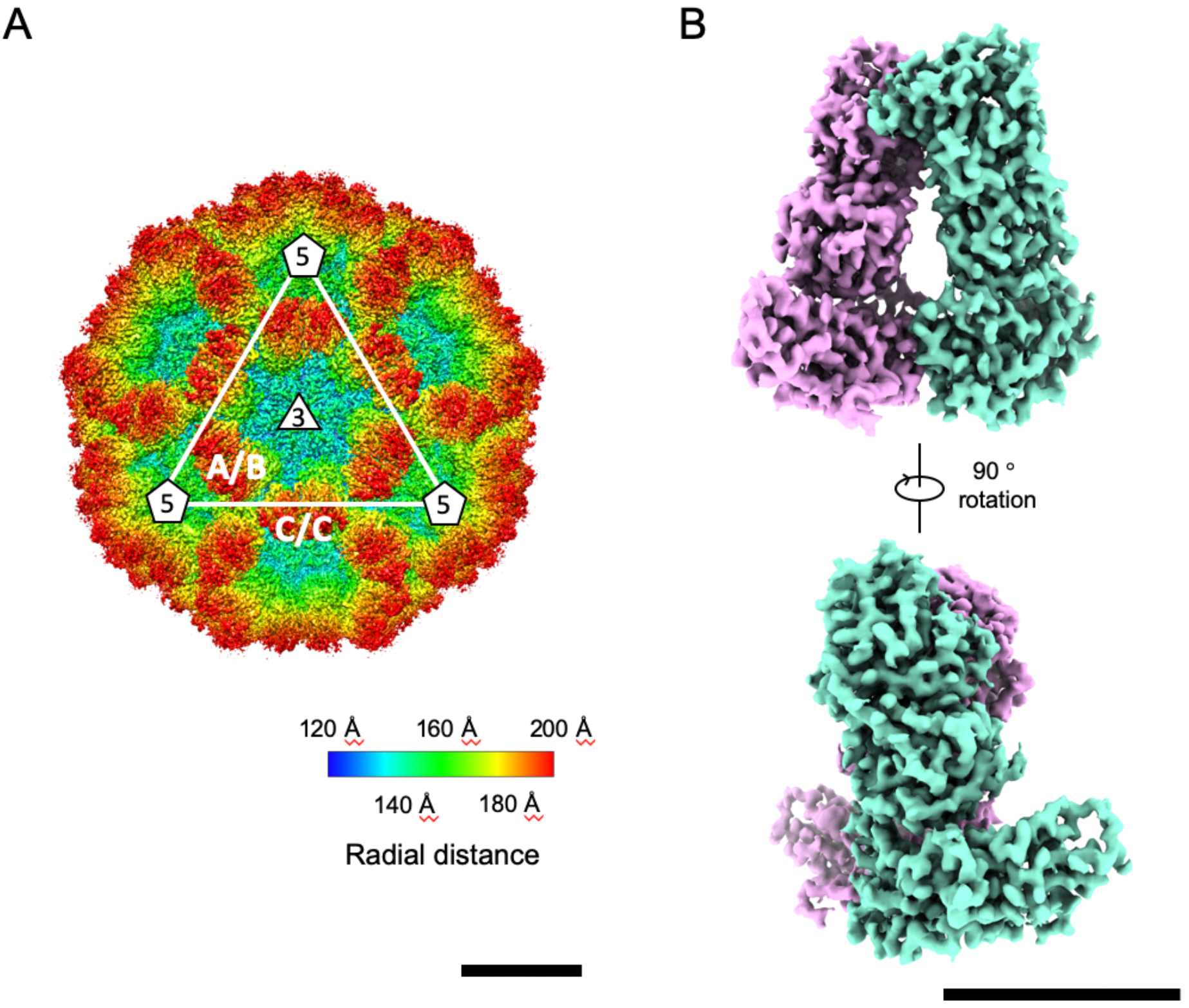
Cryo-EM maps of the HuSaV-VLP. A) Surface representation of the cryo-EM map colored according to the distance from the center of the particle. Scale bar: 10 nm. B) Surface representation of the cryo-EM map of a HuSaV VP1 C/C dimer. Monomers are colored light green and pink. Scale bar: 5 nm.

**Figure 2.**
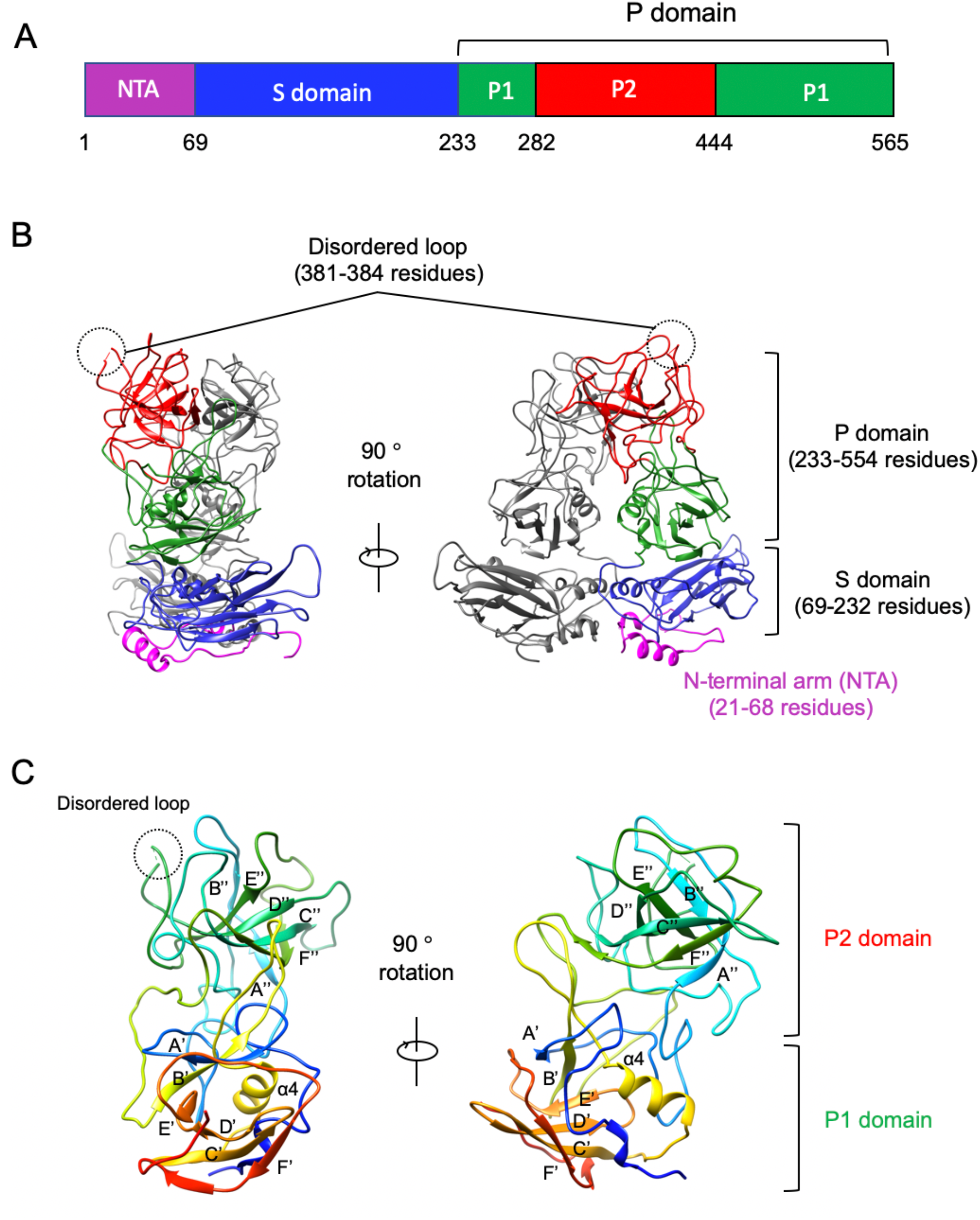
Atomic structure of the HuSaV VP1. A) Domain organization of the HuSaV VP1. N-terminal arm (NTA), S domain, P1, and P2 subdomains are colored magenta, blue, green, and red, respectively. B) Ribbon drawing of a HuSaV C/C dimer. An image (right) is drawn after the rotation of the left image by 90°. A disordered loop (residues 381–384) on the capsid surface is highlighted by a dotted circle. NTA, S domain, P1, and P2 subdomains in a monomer are colored similar to (A). C) Ribbon representation of a HuSaV P-domain. The P domain is rainbow colored with the N-terminus in blue and the C-terminus in red. An image (right) is drawn after the rotation of the left image by 90°. All secondary-structural elements are labelled.

**Figure 3.**
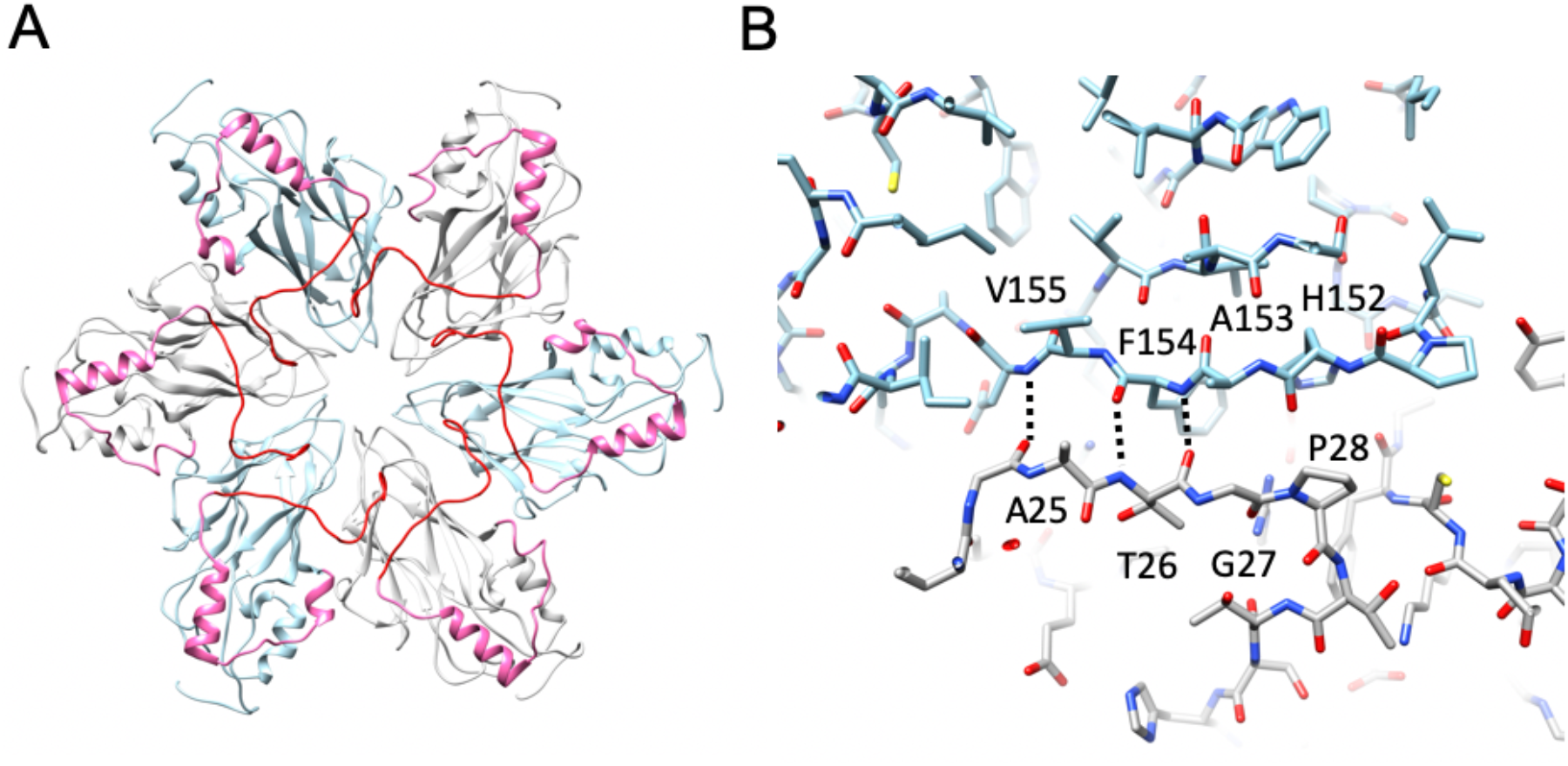
N-terminal arm network of VP1. A) Ribbon representation of S-domains around an icosahedral 3-fold axis viewed from the inside of the particle. S-domains of the B- and C-subunit are shown in gray and light blue, respectively. The N-terminal arms are highlighted by red (residues 22–37) and pink (residues 38–68). B) Inter-subunit β-sheet formed by the N-terminal arm. B- and C-subunits are shown in gray and light blue, respectively.

### The structure of the major capsid VP1 protein of HuSaV

The overall structure of the major capsid VP1 protein of HuSaV from GI. Nichinan comprises two principal domains: S (residues 69–232) and P (residues 233–554) domains, with an N-terminal arm (residues 21–68 in subunit B and C; residues 38–68 in subunit A) inside the capsid shell (Figure 2). The P domain is further divided into two subdomains, P1 (233–281 and residues 444–554) and P2 (residues 282–443) subdomains. These domain boundaries are consistent with those from previous result based on the homology model built with an 8.5-Å resolution cryo-EM map (Miyazaki et al., 2016). We found a disordered region (residues 381–384) in the P2 subdomain, which is located on the exterior surface of the viral particle (dashed circles in Figure 2B), and the βD”-βE” loop, including the disordered region, which is one of the most valuable amino acid sequences among the HuSaVs (Figure 4).

**Figure 4.**
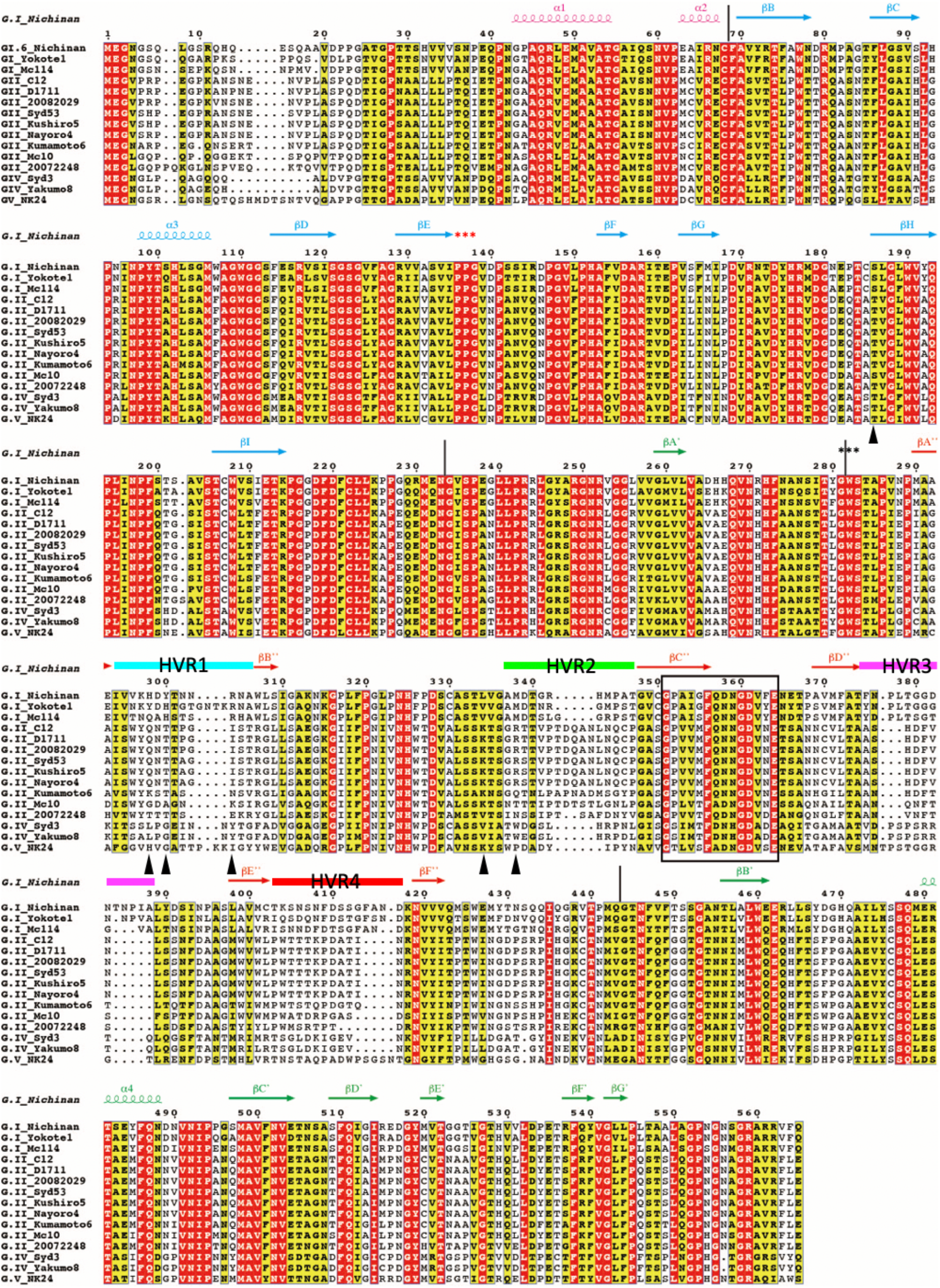
Alignment of the amino acid sequences of the HuSaV VP1 proteins. The secondary-structural elements are indicated over the sequences as a spiral (α-helix) or an arrow (β-sheet) and are colored according to the domain regions (S: pink, P1: green, P2: orange). The regions of the S, P1, and P2 domains are also shown as pink, green, and orange lines below the secondary structural elements, respectively. Letters on a red and yellow background indicate identical and similar amino acids based on a *Risler* matrix (Risler et al., 1988), respectively. Gaps are shown as dotted lines. Figure is drawn by ESPript (Gouet et al., 2003).

There are two well-conserved motifs in the SaV VP1, “PPG” (residues 136–138) and “GWS” (residues 281–283) (Oka et al, 2015), indicated by red and black asterisks above sequences in Figure 4, respectively. The former “PPG” motif exists in a βE-βF loop and is a part of β-turn (PPGV) just following a βE-strand in the S-domain, which is not exposed to the viral surface and is located at the inter-domain interface between the S-domains (Figures 5A and 5C). The loop after the β-turn interacts with the P1-subdomain (black arrowhead in Figure 5A). These observations suggest that the conserved “PPG” motif forming the β-turn is involved in protein folding, dimer formation, and assembly into the particle. The latter “GWS” motif is located at the domain boundary between the P1 and P2 subdomains (Figure 5A). The ”GWS” motif in the βA’-βA” loop is not exposed to the viral surface as in the motif “PPG” suggesting that the motif is not involved in receptor binding, particle formation, or stability because W282 in the motif is inserted into the P2 subdomain and forms a hydrophobic core with L322, I393, and M442 in the P2 subdomains (Figure 5B).

**Figure 5.**
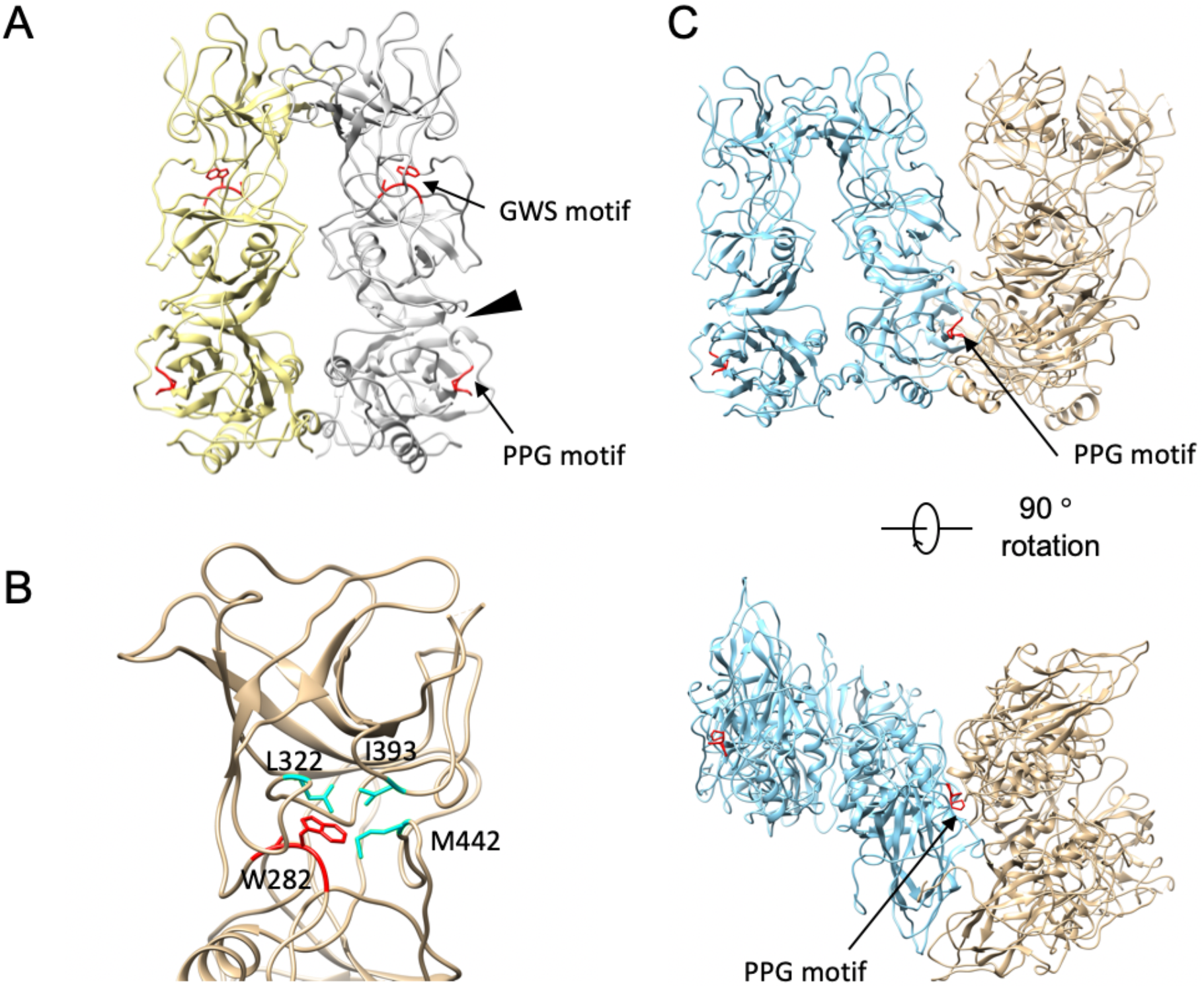
Conserved motifs among HuSaV VP1 proteins. A) Ribbon representation of a HuSaV VP1 dimer showing two well-conserved motifs in the HuSaV VP1 protein, “PPG” (residues 136–138) and “GWS” (residues 281–283), highlighted by red. An arrowhead indicates the interaction between the loop following the PPG motif and P1 subdomain. B) The “GWS” motif exists at the domain boundary between the P1 and P2 subdomains, and W282 interacts with L322, I393, and M442 in the P2 subdomain. C) The “PPG” motif, colored red, is located at the inter-subunit interface between the S-domains. A/B and C/C dimers are colored sky-blue and gold, respectively.

### Interactions between P domains in a dimeric protrusion

The P domain of HuSaV forms a dimer on the surface of virus particles, and the inter-dimer interaction occurs in the most exterior region of the dimer (Figures 1B and 6A), and the inter-dimer orientation is similar to that of SMSV and FCV (Chen et al., 2006; Ossiboff et al., 2010) as in the previous structure analysis at an 8-Å resolution (Miyazaki et al., 2016). In this study, the high-resolution structure at 2.9 Å resolution revealed the amino acid residues involved in the interaction between the P domains to build a stable P domain dimer. Unexpectedly, in addition to the residues in the P2 subdomain, the residues in the P1 subdomain are involved in the interaction between the P domains (Figure 6A). The βB’-α4 loop (residues 465–469) in the P1 subdomain and the βC”-βD” loop in the P2 subdomain are mainly involved in the interaction between the P domains. Hydrophobic residues are present throughout the binding interface, representing the hydrophobic interactions that appear to be dominant for the dimerization (Figure 6B), although some hydrophilic interactions are also observed, for example, between the βB’-α4 loop and the βC”-βD” loop. We calculated and compared buried surface areas between the P domains of caliciviruses. The buried surface area between the P domains of HuSaV is 1.1 × 10^3^ Å^2^, which is significantly smaller than those of other viruses (1.5-1.7 × 10^3^ Å^2^) (Figure 6C). These results suggest that HuSaV shows a unique arch-like dimeric protrusion in *Caliciviridae*. Furthermore, additional mechanisms may be required to stabilize the construct when the P domain dimer is used as a vaccine antigen.

**Figure 6.**
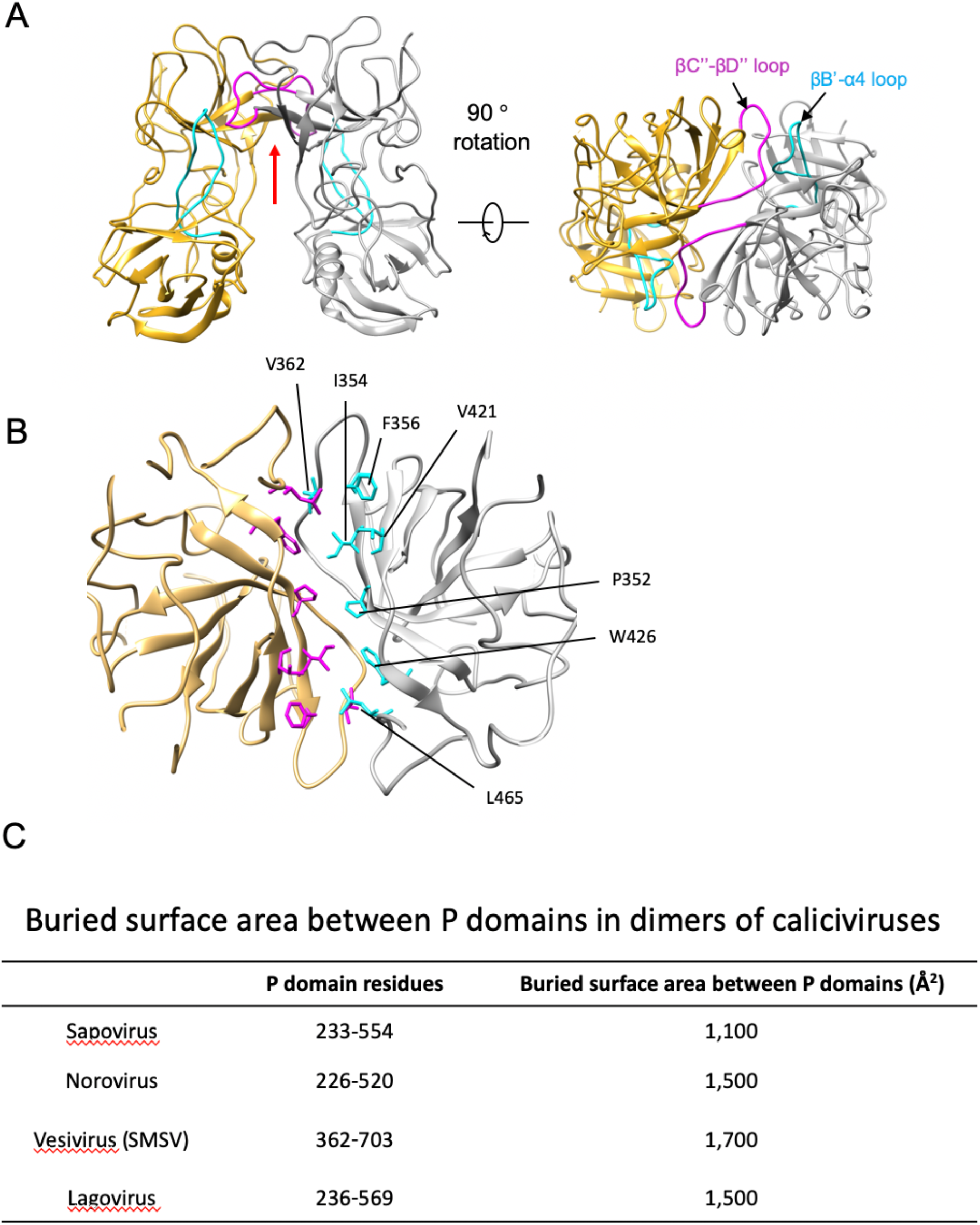
Dimeric interaction of P-domains. A) Ribbon drawing of a HuSaV P-domain dimer. Two loops, βC”-βD” and βB’-α4 loops, are highlighted by magenta and cyan, respectively. B) Hydrophobic interactions at the dimer interface viewed from the direction indicated by a red arrow in (A). C) Buried surface area between P domains in the dimers of caliciviruses.

### Immune responses of HuSaVs

To examine the immune reactivity of HuSaVs, we analyzed the conserved sequences in the structure (Figure 7). When the amino acid conservation across the human GI, GII, GIV, and GV SaVs, listed in Figure 4, are mapped on the 3D structure of VP1, the amino acid residues in the P2 subdomain on the viral surface are extremely diverged, while those in the S and P1 subdomains are highly and intermediately conserved, respectively (Figure 7A). Furthermore, the primary sequence comparison shows that there are various insertions and deletions among the HuSaV stains in the P2 subdomains (Figure 4). In particular, residues 294–306 in βA”-βB”, 337–347 in βB”-βC”, 375–388 in βD”-βE” (including a disordered region, 380–384), and 403–417 in βE”-βF” show significant sequence variabilities between the HuSaVs, which are designated as hypervariable region 1 (HVR1), HVR2, HVR3, and HVR4, respectively (Figure 4). These regions form a large cluster at the top of the P domain (Figure 7B). Generally, it is believed that the P domain projected from the viral surface is mainly involved in immune responses in caliciviruses, and the diverged amino acid sequences located on the most exterior surface are responsible for evading the host’s immune system. For instance, the neutralization antigens of FCV existing in the HVR of residues 408–529 (Matsuura Y et al., 2001; Tohya et al., 1991) are exclusively located in the P domain on the viral surface (Chen et al., 2006). HuSaV has a large cluster consisting of four HVRs on the viral surface, suggesting that the viral strategy against the host immune system, proposed by other caliciviruses, can be strongly applied to HuSaV.

**Figure 7.**
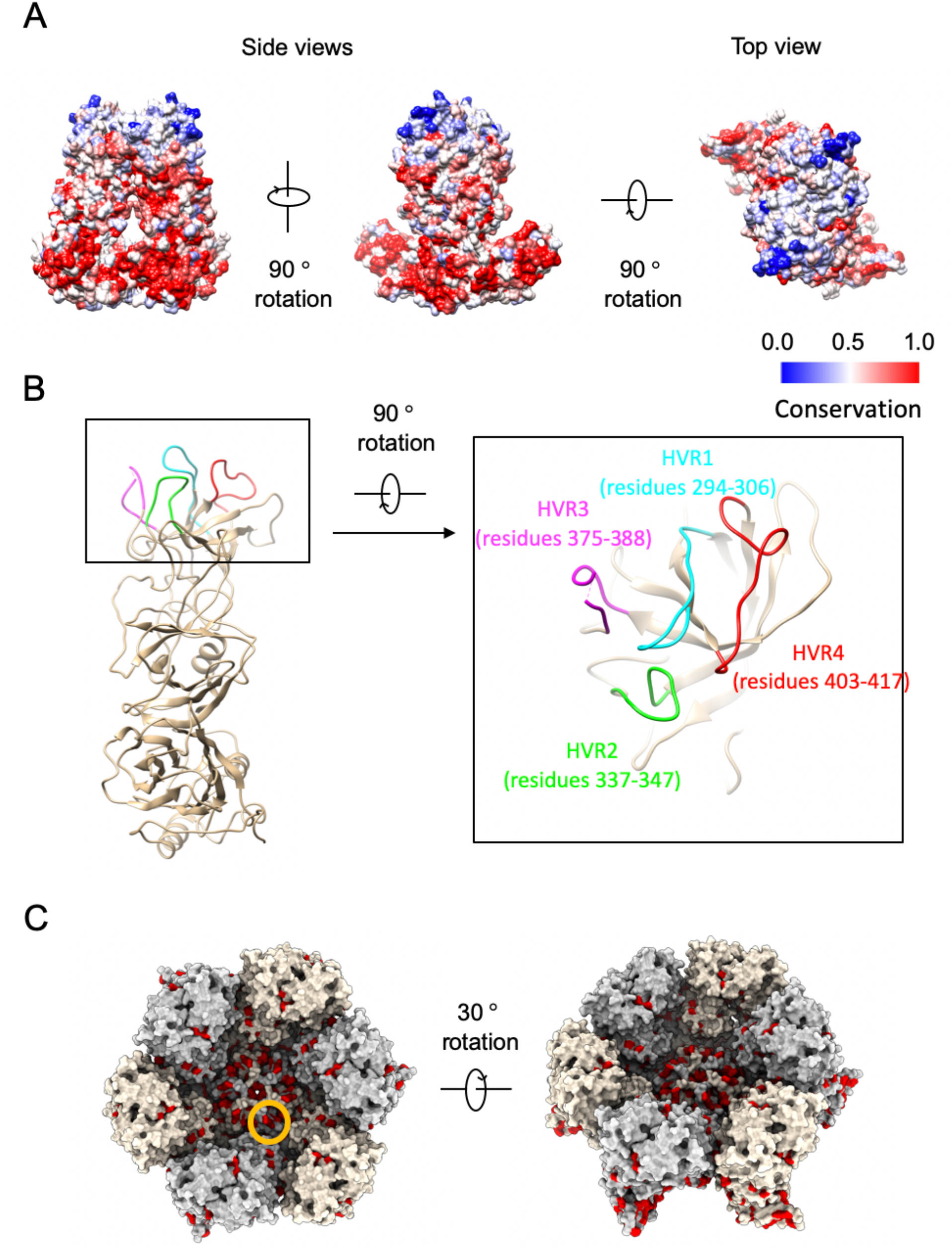
Amino acid conservation and immune-responsible surface. A) Conservation of amino acid residues of HuSaV VP1 proteins mapped onto the molecular surface of a dimer. Red indicates most conserved and blue indicates least conserved regions among the HuSaVs listed in Figure 3. B) Hyper variable regions (HVRs) of HuSaVs. Four HVRs, HVR1 to HVR4, are colored cyan, green, magenta, and red, respectively. C) Conserved amino acid residues are highlighted by red on the molecular surface of the VP1 proteins around an icosahedral 6-fold axis.

Genogroup-specific and genotype-specific monoclonal antibodies (mAbs) have been identified in a previous study (Kitamoto et al., 2012), reflecting their sequence diversity on the viral surface. However, in spite of the sequence diversity, fully cross-reactive mAbs to heterologous human genogroups and genotypes have also been found. These facts suggest that SaV capsid proteins have at least one epitope common to human GI, GII, GIV, and GV genogroups (Kitamoto et al., 2012). To examine the common epitope, we plotted the fully conserved amino acid residues (Figure 4) used in a previous immunological study (Kitamoto et al., 2012) on the molecular surface of VP1. The fully conserved amino acid residues accessible by the mAbs are limited to the S domain located in the region of the depression around the five- or six-fold axes on the viral surface (orange circle in Figure 7C). Therefore, the result suggests that one of the cross-reactive mAbs for all HuSaVs recognizes the S domain instead of the P domain.

Next, we examined epitopes recognized by mAbs specific to genogroups (GI, GII, and GIV) (Figure S4). In the case of GI and GII, the amino acid residues conserved in each genogroup form clusters in the P domain on the viral surface (magenta circles in Figures S4D-E) but not in the S domain, suggesting that the GI- and GII-specific mAbs recognize the P2 subdomain. In contrast, we found several clusters entirely in the P domain of GIV (Figures S4C and S4F). Therefore, the GIV-specific mAbs also likely bind to the P domain. However, we found clusters in the S domain (orange circle in Figure S4F), and therefore the possibility that the GIV-specific mAbs recognize the S domain cannot be completely excluded.

Genotype-specific mAbs recognize amino acid residues that are not conserved even within the genogroups. Many such amino acid residues are found in HVR1 to HVR4 on the viral surface, as described above (Figures 4 and 7A-B) as well as in the βF’’-βB’ loop. Therefore, it is considered that the highly variable regions are recognized by genotype-specific mAbs.

### Host specificity of HuSaV

The receptor molecules for HuSaV have not been identified so far, and therefore the HuSaV host recognition mechanism remains unknown. Porcine sapovirus (PoSaV) Cowden strain in a genogroup GIII is the only culturable sapovirus that has been adapted to tissue culture-adapted mutations (Lu et al., 2016; Takagi et al., 2020). Compared to the wild-type (WT) PoSaV Cowden strain, tissue culture-adapted (TC) PoSaV has six conserved amino acid substitutions in the capsid protein (Lu et al., 2016). Four of the six amino acid substitutions in VP1 (residues C178S, Y289H, M324I, and E328G) are critical for the cell culture adaptation of the PoSaV Cowden strain. Although reversion of the mutations at the other two substitutions in VP1 (residues 291 and 295) from that of the TC strain to that of the WT reduced viral replication *in vitro*, the revertants enhanced viral replication *in vivo* and induced higher-level serum and mucosal antibody responses than those induced by the TC PoSaV Cowden strain (Lu et al., 2016). We mapped these four essential and two functional mutations on the atomic structure of HuSaV. The corresponding residues for the six tissue culture-adapted mutations were S186, H298, Y300, R304, L334, and M338 (black arrowheads in Figure S5). Except for one residue S186 in the S-domain, the other five residues are exposed to the outer environment and are located on the receptor-accessible surface (Figure 8), suggesting that these residues in the P domain are actually involved in the receptor binding in the PoSaV. However, as the six residues are not conserved in HuSaV (black arrows in Figure 4), the receptor molecule of HuSaV might be different from that of PoSaV. Further studies are required to elucidate the receptor recognition mechanism of the SaV based on the atomic structure of HuSaV VP1.

**Figure 8.**
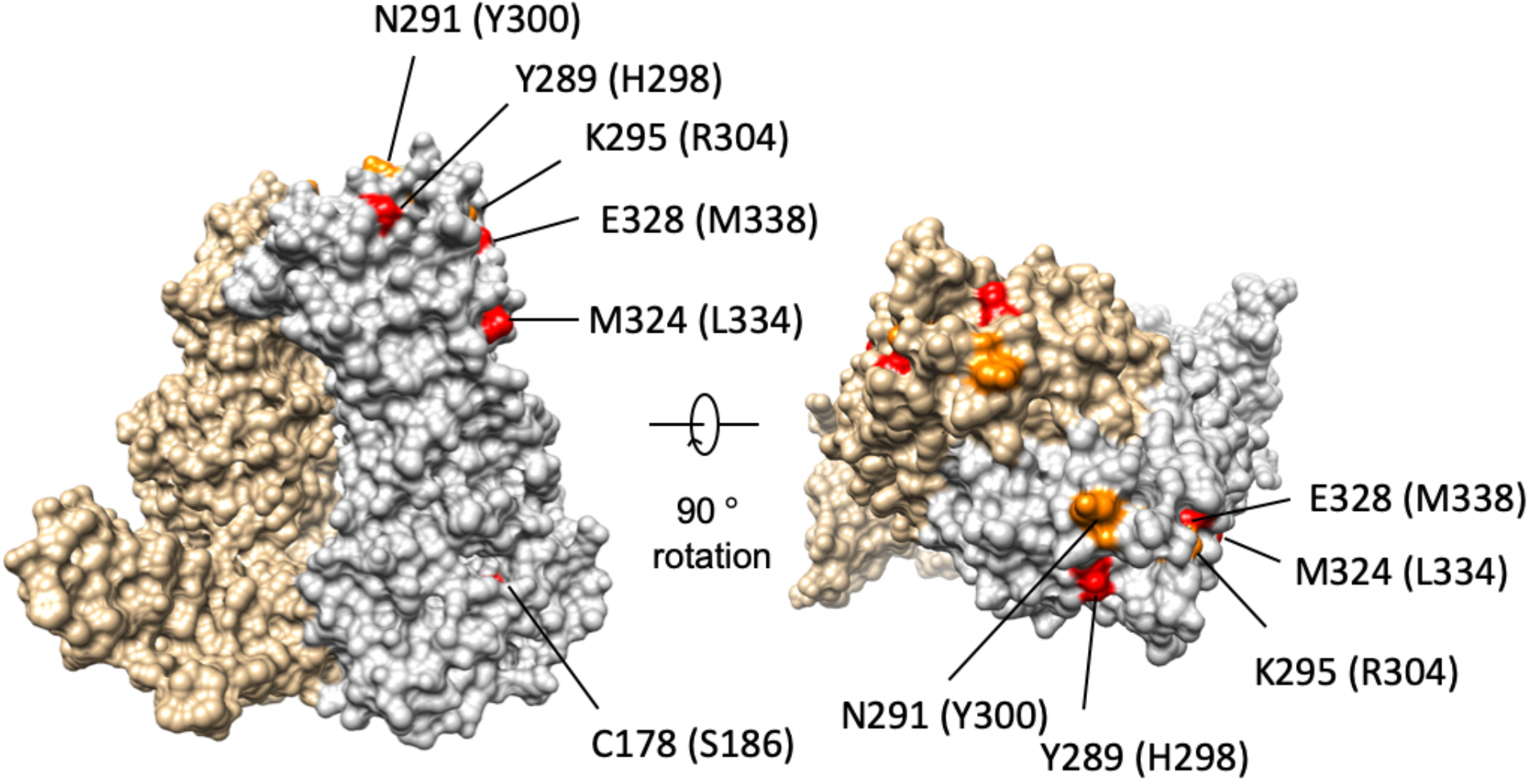
Possible receptor binding position. Six tissue culture-adapted mutation positions in the wild-type PoSaV Cowden stain are mapped on a molecular surface of a VP1 dimer in a Nichinan strain. Four essential and two functional mutation positions are labeled and colored red and orange, respectively. The corresponding amino acid residues in the Nichinan strain is enclosed in parentheses.

## Acknowledgements

This work was supported by the Platform Project for Drug Discovery, Informatics, and Structural life Science (PDIS) from the Ministry of Education, Culture, Sports, Science and Technology (MEXT), the Japan Agency for Medical Research and Development (AMED) for Supporting Drug Discovery and Life Science Research (Basis for Supporting Innovative Drug Discovery and Life Science Research (BINDS)) (Grant No. JP18am0101072 (support number 0194) to N.M. and K.I.), AMED (Grant No. 20fk0108121h0402 and 16fk0108304h2003 to Ka.M. and Grant No. 16fk0108304j9003 to K.K., Grant No. 16fk0108304j0203 to T.O.), MEXT KAKENHI (Grant No. JP16H00786 to Ka.M.), and the collaborative programs for National Institute for Physiological Sciences (to K.K.).

## Author Contributions

N.M., K.K. and Ka.M. conceived the project. T.O., Ko.M, and K.K. and M.M. expressed the VP1 protein of HuSaV and purified the HuSaV VLPs. N.M. and Ka.M. prepared cryo-EM grids and checked them at 200kV cryo-EM. N.M. and K.I. collected final high resolution cryo-EM images using 300kV cryo-EM. N.M. processed the EM data and reconstructed the final EM map. N.M. built and refined the atomic model. N.M. and Ka.M. analyzed the structure. All authors wrote the paper and contributed to experimental design and wrote the manuscript.

## Methods

### Expression of HuSaV VP1 protein in insect cells and purification of HuSaV-VLP

A baculovirus expression system constructed in a previous study (Kitamoto et al., 2012) was employed in this study. To produce VLPs of the Nichinan strain, the recombinant baculovirus of the HuSaV VP1 of the Nichinan strain was propagated with Sf9 cells (Thermo Fischer Scientific, USA) as described previously (Kitamoto et al., 2012). The recombinant baculovirus was used to infect approximately 3 × 10^6^ confluent Hi5 cells (Thermo Fischer Scientific, USA) at a multiplicity of infection (MOI) of 5–10 in 1.5 mL Ex-Cell 405 medium (Merck, Germany), and the infected cells were incubated at 26 °C. The culture medium was harvested 5–6 days post-infection (dpi), centrifuged for 10 min at 3,000 × g, and further centrifuged for 30 min at 10,000 × g. The VLPs were concentrated by ultracentrifugation for 2 h at 31,000 rpm at 4 °C (Beckman SW-31Ti rotor), and then resuspended in 500 μL of Ex-Cell 405 medium. Samples were examined for VLP formation by conventional electron microscopy after the VLPs were purified by CsCl as described previously (Miyazaki et al., 2016).

### Cryo-electron microscopy (cryo-EM) data collection and processing

For cryo-EM experiments, 3 μL of sample solution was applied to a Quantifoil holey carbon grid (R1.2/1.3, Mo 200 mesh, Quantifoil Micro Tools GmbH) at 4 °C with 100% humidity, and then plunge-frozen into liquid ethane using a Vitrobot Mark IV (Thermo Fisher Scientific, USA). The cryo-EM grids were examined at liquid nitrogen temperature using a cryo-electron microscope (Titan Krios, Thermo Fisher Scientific), incorporating a field emission gun and a Cs-corrector (Corrected electron optical systems GmbH). The microscope was operated at 300 kV and a nominal magnification of x 75,000. Movie frames were recorded using a Falcon II direct electron detector (Thermo Fisher Scientific), applied with a nominal underfocus value ranging from −1.0 to −2.5 μm. An accumulated dose of 20 electrons per Å^2^ on the sample was fractionated into a move stack of 16 image frames with 0.0625 s per frame, for a total exposure time of 1.0 s. The workflow of the cryo-EM image processes is summarized in Figure S1B. Movies (0.87 Å/pixel) were subsequently aligned and summed using MotionCorr software (Li et al., 2013) to obtain a final motion-corrected image. Estimation of the contrast transfer function was performed using the CTFFIND program (Rohou and Grigorieff, 2015). Micrographs exhibiting poor power spectra (based on the extent and regularity of the Thong rings) were rejected (4.0 Å resolution cutoff). Approximately 2,000 particles were manually picked using EMAN2 (Tang et al., 2007) and used to generate 2D classes for templates for auto-picking in Gautomatch (Zhang, 2017; K. Zhang, MRC Laboratory of Molecular Biology, Cambridge, UK, http://www.mrc-lmb.cam.ac.uk/kzhang/Gautomatch). All the following processes were performed using RELIOIN (Scheres, 2012). 79,147 auto-picked particles from 2,918 micrographs were subjected to reference-free 2D classification. A total of 77,352 particles were selected from acceptable 2D classes (Figure S1C) and were then subjected to two rounds of 3D classification with icosahedral symmetry. Finally, the 3D structure was reconstructed from 23,434 particles at 2.9 Å resolution, which was estimated by the gold standard FSC with a 0.143 cutoff (Grigorieff and Harrison, 2011). The local resolution variations were also calculated using the RELION software (Figure S2).

### Atomic model building and three-dimensional homology mapping

The 2.9 Å map was used for *de novo* atomic model construction of the VP1 protein in O (Jones et al., 1991). The initial atomic model was refined with phenix.real_space_refine (Adams et al., 2010) and manual adjustment in COOT (Emsley et al., 2010). The final model was further validated using MolProbity (Chen et al., 2010). The sequences of SaV VP1 proteins were aligned using CLUSTAL-W (Thompson et al., 1994). Identical and similar amino acid residues were defined according to the Risler matrix (Risler et al., 1988) and were mapped onto the surface of the SaV VP1 protein from GI. Nichinan using UCSF Chimera and ChimeraX software (Pettersen et al., 2004; Goddard et al., 2018).

### Data availability

The cryo-EM map of the HuSaV VLP of the Nichinan strain has been deposited in the Electron Microscopy Data Bank under accession number EMD-30793. Atomic coordinates for the atomic model of the VLP have been deposited in the Protein Data Bank under accession number 7DOD.

