## Supplementary Information for "Atomic structure of human sapovirus capsid by single particle cryo-electron microscopy"

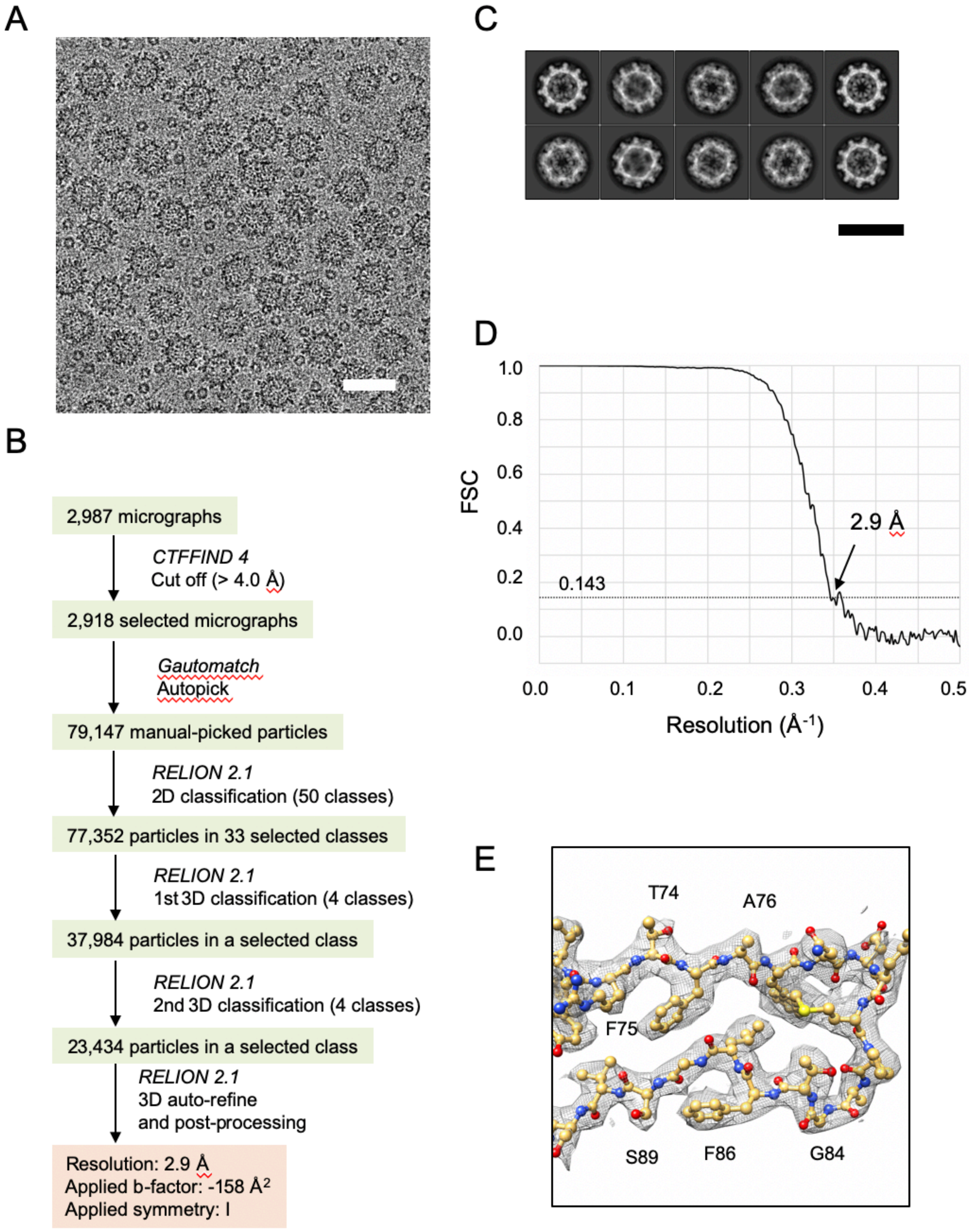

**Figure S1. Cryo-EM data collection and processing**

A) A representative cryo-EM micrograph used for image analysis of the HuSaV-VLP. Scale bar: 50 nm. B) Flowchart for the cryo-EM structure determination of the HuSaV-VLPs. C) Representative 2D class-averaged images for the HuSaV-VLPs. Scale bar: 50 nm. D) An FSC curve calculated between independently refined half maps of the reconstruction. E) Cryo-EM density map and the corresponding atomic model of the HuSaV VP1 are shown as gray meshes and ball-and-sticks, respectively. Oxygen, nitrogen, carbon, and sulfur atoms are colored red, blue, gold, and yellow, respectively.

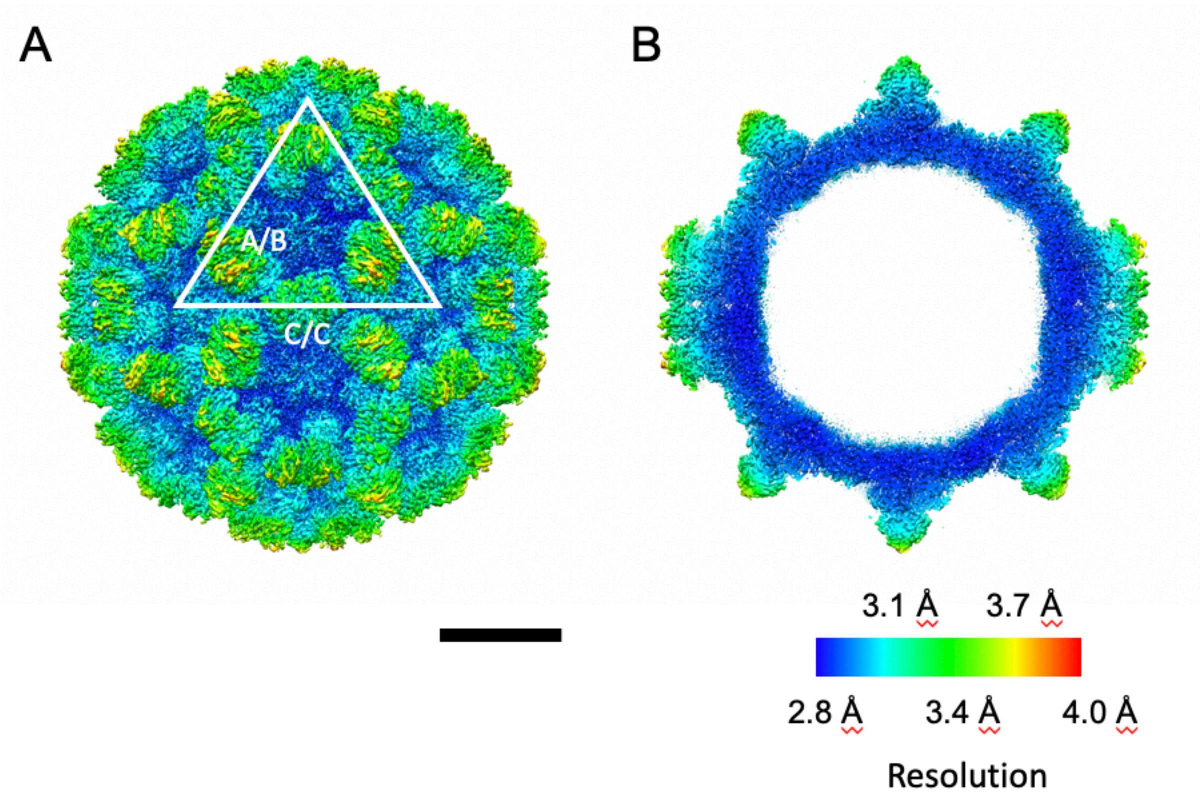


**Figure S2. Local resolution maps.** Local resolution maps of the HuSaV-VLP viewed from the outside of the particle (A) and at the central cross-section (B). Scale bar: 10 nm.

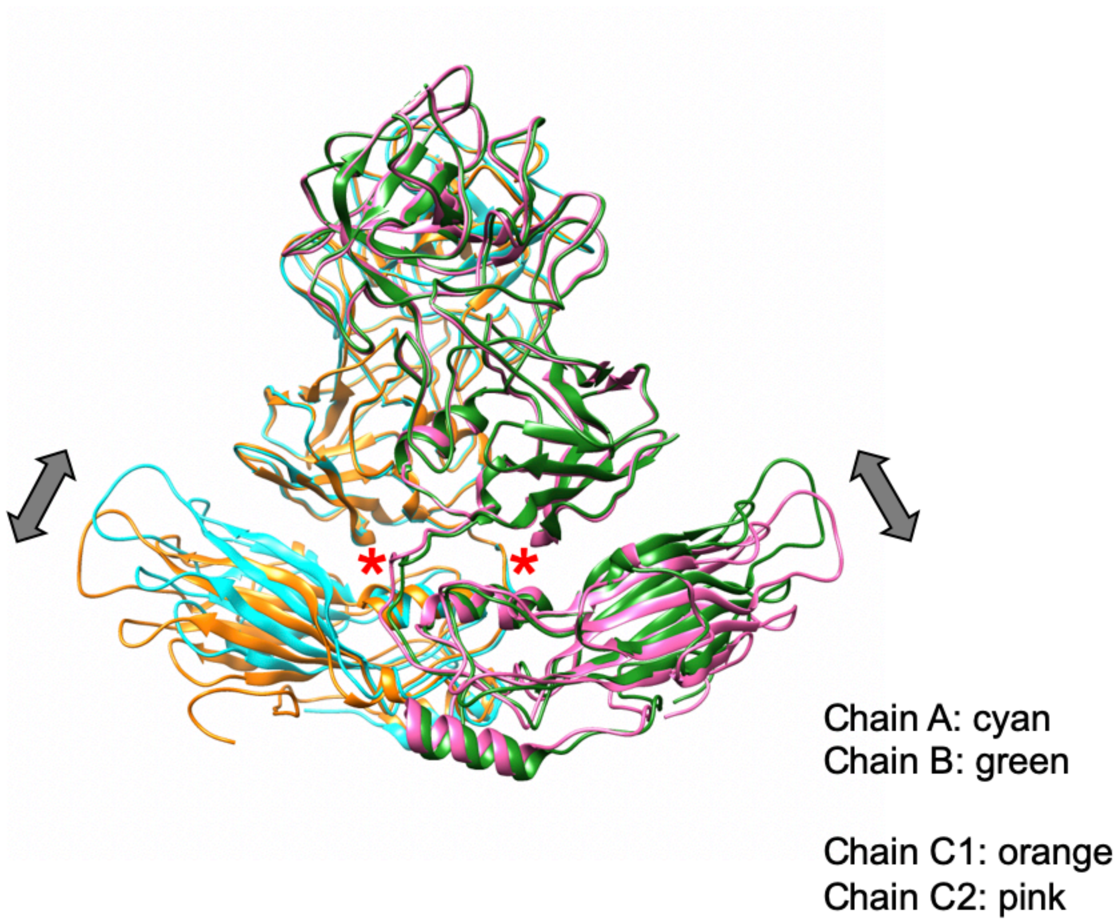


**Figure S3. Structure comparison between HuSaV A/B and C/C dimers.** Ribbon drawing of HuSaV A/B and C/C dimers colored cyan/green and orange/pink, respectively. The conformational difference is found in the S domain. The icosahedrally independent A/B and C/C dimers show slightly different conformations, which are the bent and flat conformations, respectively.

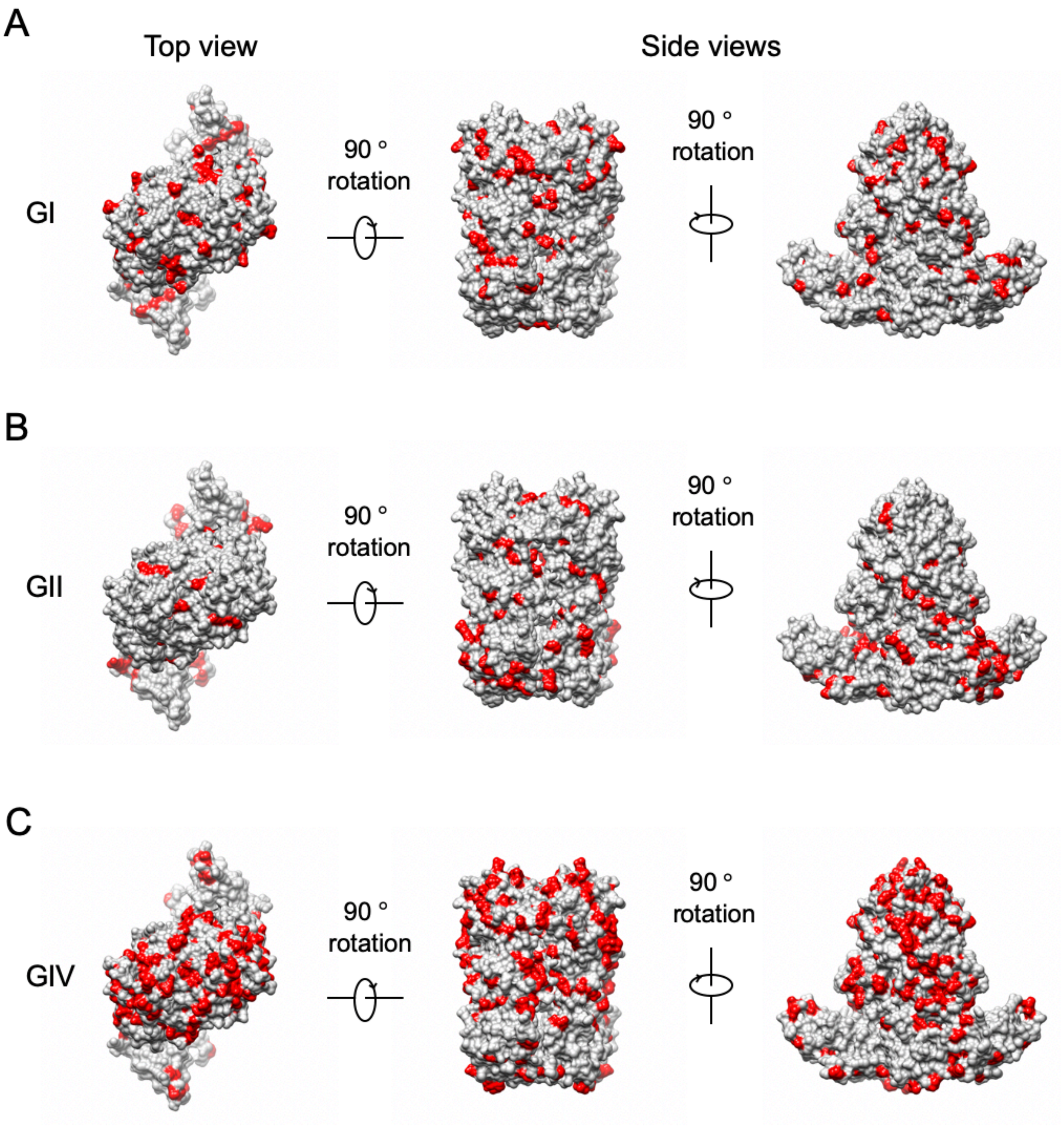
A
Top view
Side views
GI
90 °
rotation
90 °
rotation
B
GII
90 °
rotation
90 °
rotation
C
GIV
90 °
rotation
90 °
rotation

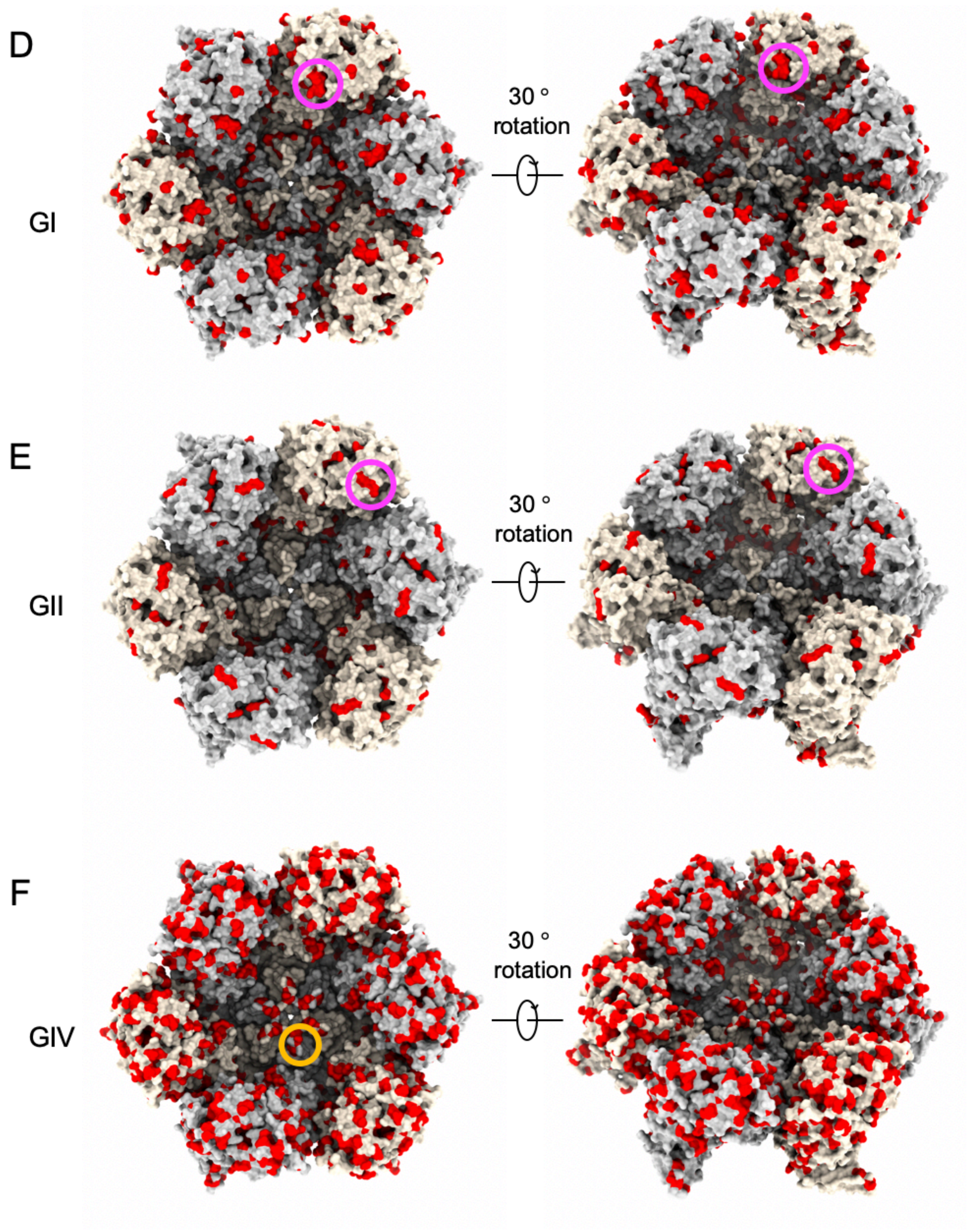
D
GI
30 °
rotation
E
GII
30 °
rotation
F
GIV
30 °
rotation

**Figure S4. Distribution of sequence conservations within genogroups of HuSaV.** Conservation of amino acid residues of HuSaV VP1 proteins mapped on the molecular surface of a dimer (A-C) and a hexamer of dimers around an icosahedral 3-fold axis viewed from the outside of the particle (D-F). The conserved amino acid residues are shown in red. A, D) Amino acid residues conserved within GI HuSaV but different from GII, GIV, and GV HuSaVs, are listed in Figure 3. B, E) Amino acid residues conserved within GII HuSaV but different from GI, GIV, and GV HuSaVs. C, F) Amino acid residues conserved within GIV HuSaV but different from GI, GII, and GV HuSaVs.

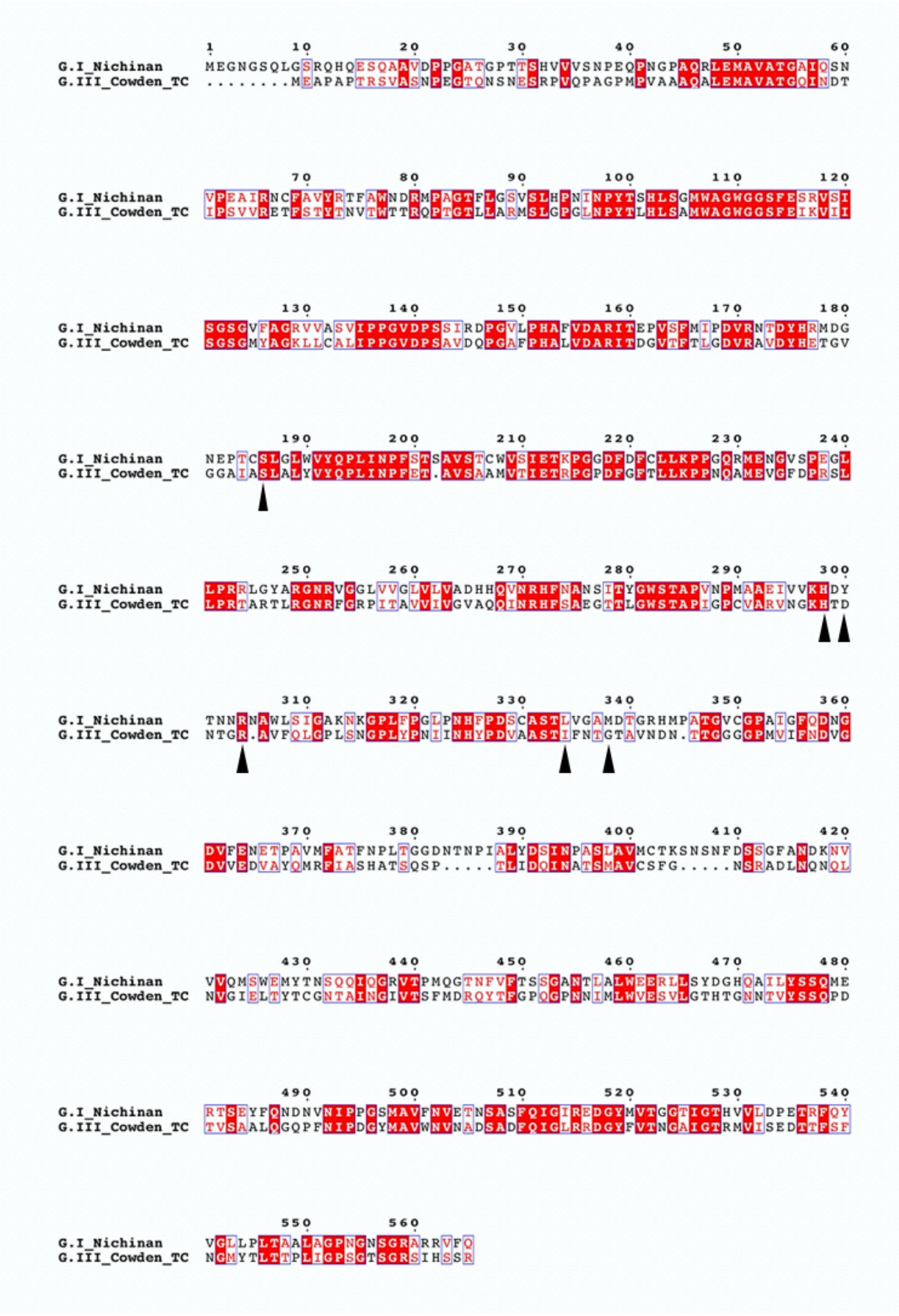
G.I_Nichinan MEGNGSQLGSRQHQESQAAVDPPGATGPTTSHVVVSNPEQPNGPAQRLEMAVATGAIQSN
G.III_Cowden_TC ........MEAPAPTRSVASNPEGTQNSNESRPVQPAGPMPVAAAQALEMAVATGQINDT
G.I_Nichinan VPEAIRNCFAVYRTFAWNDRMPAGTFLGSVSLHPNINPYTSHLSGMWAGWGGSFESRVSI
G.III_Cowden_TC IPSVVRETFSTYTNVTWTTRQPTGTLLARMSLGPGLNPYTLHLSAMWAGWGGSFEIKVII
G.I_Nichinan SGSGVFAGRVVASVIPPGVDPSSIRDPGVLPHAFVDARITEPVSFMIPDVRNTDYHRMDG
G.III_Cowden_TC SGSGMYAGKLLCALIPPGVDPSAVDQPGAFPHALVDARITDGVTFTLGDVRAVDYHETGV
G.I_Nichinan NEPTCSLGLWVYQPLINPFSTSAVSTCWVSIETKPGGDFDFCLLKPPGQRMENGVSPEGL
G.III_Cowden_TC GGAIASLALYVYQPLINPFET.AVSAAMVTIETRPGPDFGFTLLKPPNQAMEVGFDPRSL
G.I_Nichinan LPRRLGYARGNRVGGLVVGLVLVADHHQVNRHFNANSITYGWSTAPVNPMAAEIVVKHDY
G.III_Cowden_TC LPRTARTLRGNRFGRPITAVVIVGVAQQINRHFSAEGTTLGWSTAPIGPCVARVNGKHTD
G.I_Nichinan TNNRNAWLSIGAKNKGPLFPGLPNHFPDSCASTLVGAMDTGRHMPATGVCGPAIGFQDNG
G.III_Cowden_TC NTGR.AVFQLGPLSNGPLYPNIINHYPDVAASTIFNTGTAVNDN.TTGGGPMVIFNDVG
G.I_Nichinan DVFENETPAVMFATFNPLTGGDNTNPIALYDSINPASLAVMCTKSNSNFDSSGFANDKNV
G.III_Cowden_TC DVVEDVAYQMRFIASHATSQSP.....TLIDQINATSMAVCSFG.....NSRADLNQNQL
G.I_Nichinan VVQMSWEMYTNSQQIQGRVTPMQGTNFVFTSSGANTLALWEERLLSYDGHQAILYSSQME
G.III_Cowden_TC NVGIELTYTCGNTAINGIVTSFMDRQYTFGPQGPNNIMLWVESVLGTHTGNNTVYSSQPD
G.I_Nichinan RTSEYFQNDNVNIPPGSMAVFNVETNSASFQIGIREDGYMVTGGTIGTHVVLDPETRFQY
G.III_Cowden_TC TVSAALQGQPFNIPDGYMAVWNVNADSADFQIGLRRDGYFVTNGAIGTRMVISEDTTFSF
G.I_Nichinan VGLLPLTAALAGPNGNSGRARRVFQ
G.III_Cowden_TC NGMYTLTTPLIGPSGTSGRSIHSSR

**Figure S5. Alignment of the amino acid sequences between a HuSaV Nichinan strain (genogroup I) and a PoSaV Cowden strain (genogroup III)**

Letters on a red and yellow background indicate identical and similar amino acids based on a *Risler* matrix (Risler et al., 1988), respectively. Black arrowheads indicate positions for tissue culture-adapted mutations in the PoSaV Cowden strain.

Table S1. Statistics of data collection and processing

| Specimen | Sapovirus VLP |
|---|---|
| **Data collection and processing** | |
| Microscope | FEI Titan Krios G2 |
| Detector | Falcon II direct electron detector |
| Nominal magnification | 75,000 |
| Voltage (kV) | 300 |
| Nominal defocus range (μm) | −1.00 ～ −2.50 |
| Pixel size (Å) | 0.870 |
| Total electron dose ($e^-/Å^2$) | 20 |
| Exposure time (s) | 1.0 |
| Number of frames per image | 16 |
| Number of micrographs | 2,987 |
| Number of micrographs used for analysis | 2,918 |
| Initial particle number | 79,147 |
| Number of particles used for 3D reconstruction | 23,434 |
| Map resolution (Å) | 2.9 |
| Applied b-factor ($Å^2$) | -158 |
| **Model building and refinement** | |
| Refined resolution (Å) | 2.9 |
| Map CC (around atoms) | 0.795 |
| Rmsd bond length (Å) | 0.01 |
| Rmsd bond angles (°) | 1.00 |
| Ramachandran plot preferred (%) | 5.64 |
| Ramachandran plot allowed (%) | 94.36 |
| Ramachandran plot outlier (%) | 0.0 |
| Rotamer outlier (%) | 0.15 |
| Clashscore | 6.68 |
| MolProbity score | 1.75 |
